# Insights into Gastrointestinal Redox Dysregulation in a Rat Model of Alzheimer’s Disease and the Assessment of the Protective Potential of D-galactose

**DOI:** 10.1101/2023.07.27.550831

**Authors:** Jan Homolak, Konstantinos Varvaras, Vittorio Sciacca, Ana Babic Perhoc, Davor Virag, Ana Knezovic, Jelena Osmanovic Barilar, Melita Salkovic-Petrisic

## Abstract

**Introduction:** Recent evidence suggests that the gut plays a vital role in the development and progression of Alzheimer’s disease (AD) by triggering systemic inflammation and oxidative stress. The well-established rat model of AD, induced by intracerebroventricular administration of streptozotocin (STZ-icv), provides valuable insights into the GI implications of neurodegeneration. Notably, this model leads to pathophysiological changes in the gut, including redox dyshomeostasis, resulting from central neuropathology. Our study aimed to investigate the mechanisms underlying gut redox dyshomeostasis and assess the effects of D-galactose, which is known to benefit gut redox homeostasis and alleviate cognitive deficits in this model.

**Materials and Methods:** Duodenal rings isolated from STZ-icv animals and control groups were subjected to a prooxidative environment using AAPH or H_2_O_2_ with or without D-galactose in oxygenated Krebs buffer ex vivo. Redox homeostasis was analyzed through protein microarrays and functional biochemical assays, alongside cell survival assessment. Structural equation modeling, univariate, and multivariate models were employed to evaluate the differential response of STZ-icv and control samples to the controlled prooxidative challenge.

**Results:** STZ-icv samples showed suppressed expression of catalase and glutathione peroxidase 4 (GPX4) and increased baseline activity of enzymes involved in H_2_O_2_ and superoxide homeostasis. The altered redox homeostasis status was associated with an inability to respond to oxidative challenges and D-galactose. Conversely, the presence of D-galactose increased antioxidant capacity, enhanced catalase and peroxidase activity, and upregulated superoxide dismutases in the control samples.

**Conclusion:** STZ-icv-induced gut dysfunction is characterized by a diminished ability of the redox regulatory system to maintain long-term protection through the transcription of antioxidant response genes, as well as compromised activation of enzymes responsible for immediate antioxidant defense. D-galactose can exert beneficial effects on gut redox homeostasis under physiological conditions.

## Introduction

A growing body of evidence suggests that bidirectional communication between the gut and the brain may play an important role in the development and progression of neurodegenerative disorders, including Alzheimer’s disease (AD)[1]. The gastrointestinal (GI) tract serves as a potential gateway for toxins, amyloidogenic proteins and microorganisms due to its large surface area, vast number of intraluminal microbes, and the significant vulnerability of the GI barrier [2,3]. Moreover, the gut acts as the body’s largest immune organ, capable of initiating and maintaining systemic inflammation and oxidative stress, which can have significant impacts on overall organismic homeostasis[4–6]. As a result, these processes can lead to neuroinflammation, insulin resistance, and ultimately neurodegeneration[7–12]. The communication from the brain to the gut plays a vital role in preserving the structural and functional integrity of the GI barrier[5,13,14]. As a result, even when neurodegeneration primarily affects the central nervous system (CNS), the GI barrier’s integrity will inevitably be compromised, leading to a harmful pathophysiological positive feedback loop between the brain and the gut. Understanding the pathophysiological changes in the GI tract related to neuropathological changes in the brain, is a crucial area of research that offers an opportunity to slow down the progression of neurodegenerative disorders by preventing chronic inflammation and oxidative stress caused by altered GI homeostasis and the breakdown of the gut-brain axis.

In transgenic animal models of AD, it is common to observe physiological changes in the GI tract before observing any pathological changes in the brain[15–17]. This sequence of events makes it difficult to study the GI consequences of localized neurodegeneration in the CNS. On the other hand, brain-first non-transgenic animal models, which involve the targeted delivery of toxins that mimic AD-related neuropathology to the CNS, provide an opportunity to investigate the one-way disruption of the gut-brain axis [18–21]. These models offer invaluable insights into how localized neurodegeneration impacts the structural and functional integrity of the gut. One of the most extensively studied and well-established brain-first models of sporadic AD is the rat model generated by intracerebroventricular administration of the diabetogenic toxin streptozotocin (STZ-icv)[22,23]. Based on the evidence accumulated over the last 30 years, the STZ-icv model recapitulates key behavioral and neuropathological hallmarks of AD[22]. Following toxin administration, the animals develop insulin resistant brain state associated with mitochondrial dysfunction and glucose hypometabolism[24,25], oxidative stress[26], neuroinflammation[27,28], neuropathological changes related to accumulation of amyloid β and hyperphosphorylated tau[29,30], chronic and progressive cognitive deficits and circadian dysrhythmia[22,31,32]. Interestingly, dysfunctional gut-brain axis and pathophysiological changes in the gut were also reported in this model indicating it represents a valid platform for investigating the GI consequences of neurodegeneration affecting the CNS. The STZ-icv gut is characterized by violated structural and functional properties of the GI barrier[33], distorted secretion and constitution of gut mucus[20], redox dyshomeostasis[18,34], and altered absorption[35].

Given the significant role of redox homeostasis in maintaining gut function and GI barrier integrity[4], the main focus of this study was to employ an *ex vivo* approach to investigate the underlying mechanisms behind the observed redox dyshomeostasis in the gut of the STZ-icv model. The aim was to subject intestinal rings to a controlled oxidative environment and assess their ability to withstand an exogenous oxidative challenge, which necessitates functional redox homeostasis. The secondary goal of the study was to investigate the effects of D-galactose on redox homeostasis in the STZ-icv gut. This was motivated by previous findings that while parenteral D-galactose administration has been linked to oxidative stress and cognitive decline[36], chronic oral D-galactose treatment has shown promising results in preventing and alleviating cognitive deficits in the STZ-icv model[25,37–39]. Additionally, orally administered D-galactose has been found to have beneficial effects on redox homeostasis both in the GI tract[40] and the brain[41].

## Materials and methods

### Rat model of sporadic Alzheimer’s disease

The Croatian Ministry of Agriculture (EP 186 /2018) and the Ethical Committee of the University of Zagreb School of Medicine (380-59-10106-18-111/173) approved all procedures. To establish the rat model of sAD, we utilized the intracerebroventricular administration of streptozotocin (STZ-icv) following a standard procedure [18,22,23,27]. A total of eight 12-weeks-old male Wistar rats, bred under standardized conditions at the Department of Pharmacology, were randomly divided into two groups. The rats were anesthetized using intraperitoneal administration of ketamine/xylazine (70/7 mg/kg) and then subjected to intracerebroventricular treatment at specific coordinates relative to bregma (−1.5 mm posterior; ± 1.5 mm lateral; +4 mm ventral) [42]. Streptozotocin (STZ) was freshly dissolved in 0.05 M citrate buffer (pH 4.5). Control animals (CTR; n=4) were bilaterally injected with vehicle (2 μl/ventricle) and rats used for modelling sAD (STZ; n=4) received STZ solution (2 μl/ventricle; 1.5 mg/kg). The same procedure was repeated after 48 hours to achieve a cumulative dose of 3 mg/kg of STZ [22,23,27].

### Tissue harvesting for *ex vivo* experiments

Duodenal rings for *ex vivo* experiments were obtained from 20-weeks-old animals (2 months after STZ-icv). The animals were anesthetized (i.p. ketamine/xylazine; 70/7 mg/kg) and decapitated. Proximal duodenum was isolated and washed with pre-heated (35 °C) Krebs buffer (115 mM NaCl, 25 mM NaHCO_3_, 2.4 mM K_2_HPO_4_, 1.2 mM CaCl_2_, 1.2 mM MgCl_2_, 0.4 mM KH_2_PO_4_, 10 mM glucose bubbled with Carbogen gas (95% O_2_; 5% CO_2_) to remove intraluminal contents. Tissue was placed on top of a wet cellulose paper towel in a Petri dish filled with Krebs buffer and cut into 4 mm thick duodenal rings.

### Treatments

Duodenal rings were incubated in a 96 well plate for 30 minutes. Control sections were incubated in Krebs buffer (CTR); pro-oxidative environment was modelled by incubation with 200 μM 2,2’-Azobis(2-amidinopropane) dihydrochloride in Krebs buffer (AAPH) or 1.5 mM H_2_O_2_ in Krebs buffer (H2O2). The ability of D-galactose to protect the tissue against oxidative challenge was investigated by incubation with 100 mM D-galactose in the presence of 200 μM AAPH (AAPH+GAL) or 1.5 mM H_2_O_2_ (H2O2+GAL).

### Tissue preparation

Duodenal rings were removed form the experimental solution, rinsed in Krebs buffer, snap-frozen in liquid nitrogen and stored at −80°C. Samples were homogenized with ultrasonic waves (Microson Ultrasonic Cell 167 Disruptor XL, Misonix, Farmingdale, NY, SAD) in lysis buffer containing 150 mM NaCl, 50 mM Tris-HCl, 1 mM EDTA, 1% Triton X-100, 1% sodium deoxycholate, 0.1% SDS, 1 mM PMSF, protease (Sigma-Aldrich, Burlington, MA, USA) and phosphatase inhibitor (PhosSTOP, Roche, Basel, Switzerland) cocktail (pH 7.5). Homogenates were centrifuged (relative centrifugal force (RCF) 12 879 g) for 10 min at 4 °C. Protein concentration in supernatants was analyzed with the Bradford reagent (Sigma-Aldrich, USA) using bovine serum albumin in lysis buffer as a standard solution. Samples were stored at −80°C.

### DAPI permeability assay

DAPI permeability assays was performed to label dead cells for subsequent analysis. A set of duodenal rings adjacent to the rings used for the analysis of redox biomarkers and subjected to the same treatment (CTR, AAPH, H2O2, AAPH+GAL, H2O2+GAL) was rinsed in Krebs buffer and immersed for 5 min in Krebs buffer containing 4′,6-diamidino-2-phenylindole (DAPI; 1 μg/ml). After incubation samples were rinsed in Krebs buffer and stored in 4% paraformaldehyde (pH 7.4) at 4 °C. Tissue sections were cut using a cryostat (Leica, Germany) following fixation in Tissue-Tek O.C.T (Akura Finetek USA, USA). Samples were coverslipped with fluoroshield mounting medium (9 parts glycerol, 1 part 10XPBS, 2% n-propyl gallate) and analyzed using the U-MNU2 filter set (EX: 365/10; EM: >420) on the Olympus BX51 epifluorescent microscope (Olympus, Japan). Images were analyzed in Fiji (NIH, USA).

### Analysis of redox biomarkers

Total antioxidant capacity was analyzed with 2,2′-azino-bis(3-ethylbenzothiazoline-6-sulfonic acid)(ABTS) radical cation assay[43] and nitrocellulose redox permanganometry (NRP)[44]. Briefly, ABTS radical cation was generated by reacting 7 mM ABTS with 2.45 mM K_2_S_2_O_8_ in dark for 24 hours. The solution was diluted 40-fold in PBS to obtain optimal baseline absorbance. Tissue homogenates (1 μl) were incubated with 100 μl of the ABTS working solution in 96-well plates and 405 nm absorbance was recorded after 5 min using an Infinite F200 PRO multimodal microplate reader (Tecan, Switzerland). 1,4-dithiothreitol (DTT) was used as a standard reducing agent[45]. NRP was measured by first spotting 1 μl of each sample on the nitrocellulose membrane (Amersham Protran 0.45; GE Healthcare Life Sciences, USA). Dry membrane was immersed in NRP working solution (10mg/ml KMnO_4_ in ddH_2_O). The membrane was destained in dH_2_O, dried, scanned and analyzed in Fiji with Gel Analyzer plugin as described previously[44]. Activity of SODs was determined with a modified 1,2,3-trihydroxybenzene (THB) autoxidation inhibition assay [46–48]. Reaction buffer contained 0.05 M Tris-HCl and 1 mM Na_2_EDTA (pH 8.2) and the final concentration of THB was 1.2 mM. Reaction buffer was modified by adding 2 mM KCN to discriminate between Cu/Zn- and Mn/Fe-SOD[21]. THB autoxidation was monitored at 450 nm with a multimodal microplate reader (Tecan, Switzerland). Catalase and peroxidase activities were determined indirectly by quantifying residual H_2_O_2_ assessed by Co oxidation after incubation with 10 mM H_2_O_2_ in PBS[49]. Samples (10 μl) were first incubated with 40 μl of 10 mM H_2_O_2_ for 90 s and then reacted with 100 μl of Co(NO_3_)_2_ working solution. Baseline interference was ruled out by reacting the samples with stop solution before adding the substrate[21,50]. Carbonato-cobaltate (III) complex ([Co(CO_3_)_3_]Co) absorbance was measured at 450 nm with a microplate reader (Tecan, Switzerland) and compared to the H_2_O_2_ standard model to determine H_2_O_2_ concentration. The same procedure was repeated in the presence of 25 μM NaN_3_ to discriminate between the activity of catalase and peroxidases[21,51]. Reaction conditions were optimzied in pilot experiments. Lipid peroxidation was assessed with the thiobarbituric acid reactive substances (TBARS) assay[41]. Tissue homogenates (20 μl) were mixed with the TBA-TCA solution (0.4% thiobarbituric acid (Kemika, Croatia); 15% trichloroacetic acid (Sigma-Aldrich, USA)) in perforated microcentrifuge tubes. The samples were boiled at 95 °C for 45 min and the colored adduct was extracted with n-butanol. The absorbance of butanol extracts was measured at 540 nm in a 384-well plate using the Infinite F200 PRO multimodal plate reader (Tecan, Switzerland). Malondialdehyde tetrabutylammonium in ddH_2_O (Sigma-Aldrich, USA) serial dilutions were processed in parallel to obtain a standard curve. Reaction steps were optimized in pilot experiments.

### Protein microarray

Expression of proteins of interest (catalase (CAT), superoxide dismutase 1 (SOD1), superoxide dismutase 2 (SOD2), glutathione peroxidase 4 (GPX4), and neuronal nitric oxide synthase (nNOS)) was analyzed with indirect immunofluorescence-based microarray. Samples were fixed onto a nitrocellulose membrane using a PVC template. Fixed samples were blocked in a blocking buffer (10 mM Tris, 150 mM NaCl, 5% (w/v) nonfat dry milk, 0,5% (v/v) Tween 20 (pH 7.5)) at room temperature for 1 hour. Membranes were incubated with primary antibodies diluted in the blocking buffer (anti-CAT (ABIN872991; 1:400), anti-SOD1(ABIN2854826; 1:400), anti-SOD2 (PA1776; 1:400), anti-GPX4 (CQA1094; 1:400), anti-nNOS (PA1329; 1:400)) for 24 hours at +4 °C. Membranes were washed in 1xPBS and incubated with anti-rabbit IgG (H+L) F(ab’)_2_ fragment conjugated to Alexa Fluor® 488 (4412S; 1:500; Cell Signaling Technology, USA) for 2 hours at room temperature. Unbound antibodies were removed in 1xPBS and sample fluorescence was recorded using ChemiDoc MP Imaging System (Bio-Rad, USA)(EX/EM (nm): 460-490/518-546). Background fluorescence was recorded (EX/EM (nm): 302/535-645) for adjustment. Protein expression data was obtained using a pipeline in which expression was determined based on 460-490/518-546 fluorescence adjusted for autofluorescence (302/535-645) and protein concentration.

### Quantification of nitrites

The Griess assay was utilized to indirectly quantify nitric oxide (NO) by measuring the nitrite concentration. The modified Griess reagent (Sigma-Aldrich, USA), consisting of naphthylethylenediamine dihydrochloride and sulphanilamide in phosphoric acid, was employed in the assay. To accommodate the 384-well plate format, optimal sample volumes were adjusted and mixed in a 1:1 volumetric ratio with the Griess working solution (0.4% (w/v)). The absorbance change was measured at 540 nm over a period of 15 minutes.

### Data analysis

Data analysis was was conducted using R (4.1.3) adhering to the guidelines for reporting animal experiments[52]. The statistical modeling approach employed linear-mixed models to address the hierarchical structure of the data. In this approach, redox biomarkers were designated as the dependent variables, while the independent variables included group (CTR vs. STZ), treatment (CTR, AAPH, H2O2, AAPH+GAL, H2O2+GAL), and the interaction between group and treatment. Moreover, protein concentration was included as a covariate to address any potential bias that may have arisen during sample preparation procedures. Additional covariates were also introduced when necessary, such as baseline H_2_O_2_ in the CAT model, to further control for relevant factors. Sample non-specific fluorescence was introduced as an additional covariate in protein expression models. Animal identification number was defined as a random effect to account for hierarchical structure. Model assumptions were verified through visual inspection of residual and fitted value plots, and if required, variable transformations were applied. Model estimates were presented as effect sizes along with their corresponding 95% confidence intervals. H_2_O_2_ and SOD signaling pathways were modeled with structural equation modeling (SEM) in *lavaan*[53]. Enzyme expression and activity data were utilized as the manifest variables to create latent variables for H_2_O_2_ and SOD signaling. In the H_2_O_2_ model, the covariance between CAT expression and baseline H_2_O_2_, as well as the covariance between total H_2_O_2_ dissociation capacity and baseline H_2_O_2_, were both included. Similarly, in the SOD model, the covariance between SOD1 and SOD2 expression, and the covariance between SOD2 expression and SOD2 activity were incorporated based on modification indices. To assess the suitability of SEMs, goodness-of-fit indices such as Comparative Fit Index (CFI) and Tucker-Lewis Index (TLI) and badness-of-fit indices including Root Mean Square Error of Approximation (RMSEA) and Standardized Root Mean Square Residual (SRMSR) were evaluated. Standardized loadings were reported to assess the relationships between latent and manifest variables. Differences of H_2_O_2_ and SOD pathways between groups (CTR vs. STZ) were analyzed with a χ^2^ difference test. α was fixed at 5%. Nonlinear dimensionality reduction was performed using uniform manifold approximation and projection (UMAP). Both SEM and UMAP analyses were performed on scaled datasets.

## Results

To evaluate the impact of prooxidative environments (AAPH, H2O2) and the potential protective effects of D-galactose on both CTR and STZ intestinal cells, we initiated our analysis by examining the survival of mucosal cells (Fig 1A, B). The number of dead cells under normal conditions (Krebs buffer) was comparable between the control group (CTR) and the STZ-treated group (Fig 2C). The only instance where a noticeable distinction in cell survival between the CTR and STZ tissue was observed was during incubation with H_2_O_2_. This resulted in reduced estimates of dying cells in the CTR group and decreased survival in the STZ mucosa (Fig 1C). The diminished survival of STZ cells was partially mitigated when co-incubated with D-galactose; however, it is important to note that these effects were accompanied by significant uncertainty (Fig 1C). In STZ tissue, the biomarker NRP, which indicates total antioxidant capacity[44], showed no significant response. However, in the control group, the NRP levels were higher in the prooxidative environment regardless of the presence of D-galactose (Fig 1D). Interestingly, the ABTS assay indicated a reduced overall antioxidant capacity in the STZ tissue, which was not improved by co-incubation with D-galactose (Fig 1E). It is worth noting that, except for the increased antioxidant capacity (NRP) observed in the control tissues incubated with D-galactose in the presence of AAPH, both biomarkers of total antioxidant capacity indicated that the effects were modest and accompanied by considerable uncertainty estimates (Fig 1 D,E).

**Fig 1.**
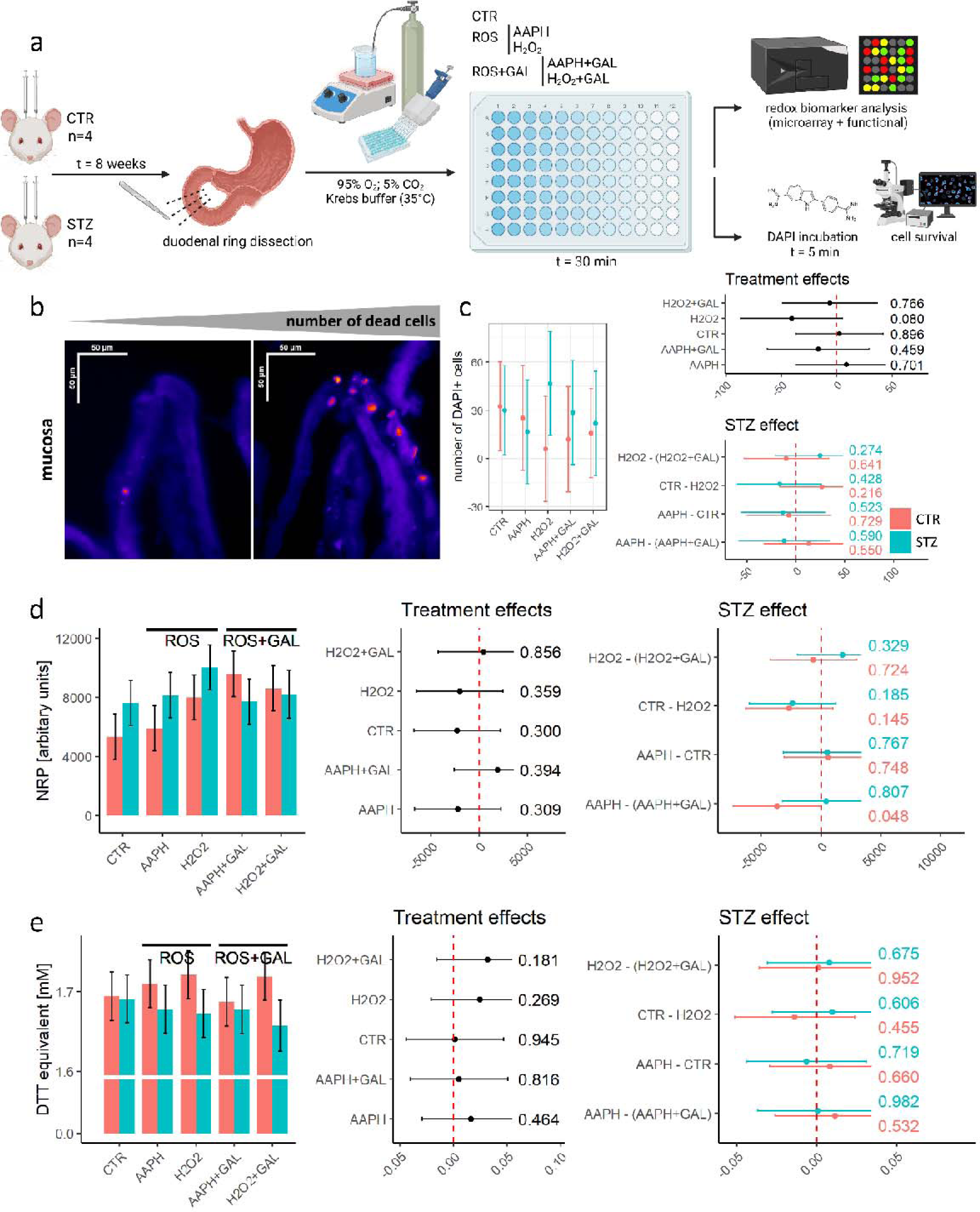
Experimental design and the analysis of cell survival and antioxidant capacity. Experimental design (a); Representative photomicrograph of DAPI signal with dead cells shown in warm colors (*Fire* Look-Up Table)(b); quantification of dead cells with group estimates (left) and effect sizes (right)(c); NRP signal group estimates (left) and effect sizes (right)(d). ABTS signal group estimates (left) and effect sizes (right)(e). The error bars in the bar graphs represent standard errors, while for effect size estimates, 95% confidence intervals are provided. P-values are reported alongside the model-derived estimates. ROS – reactive oxygen species (prooxidative environment); GAL – D-galactose.

**Fig 2.**
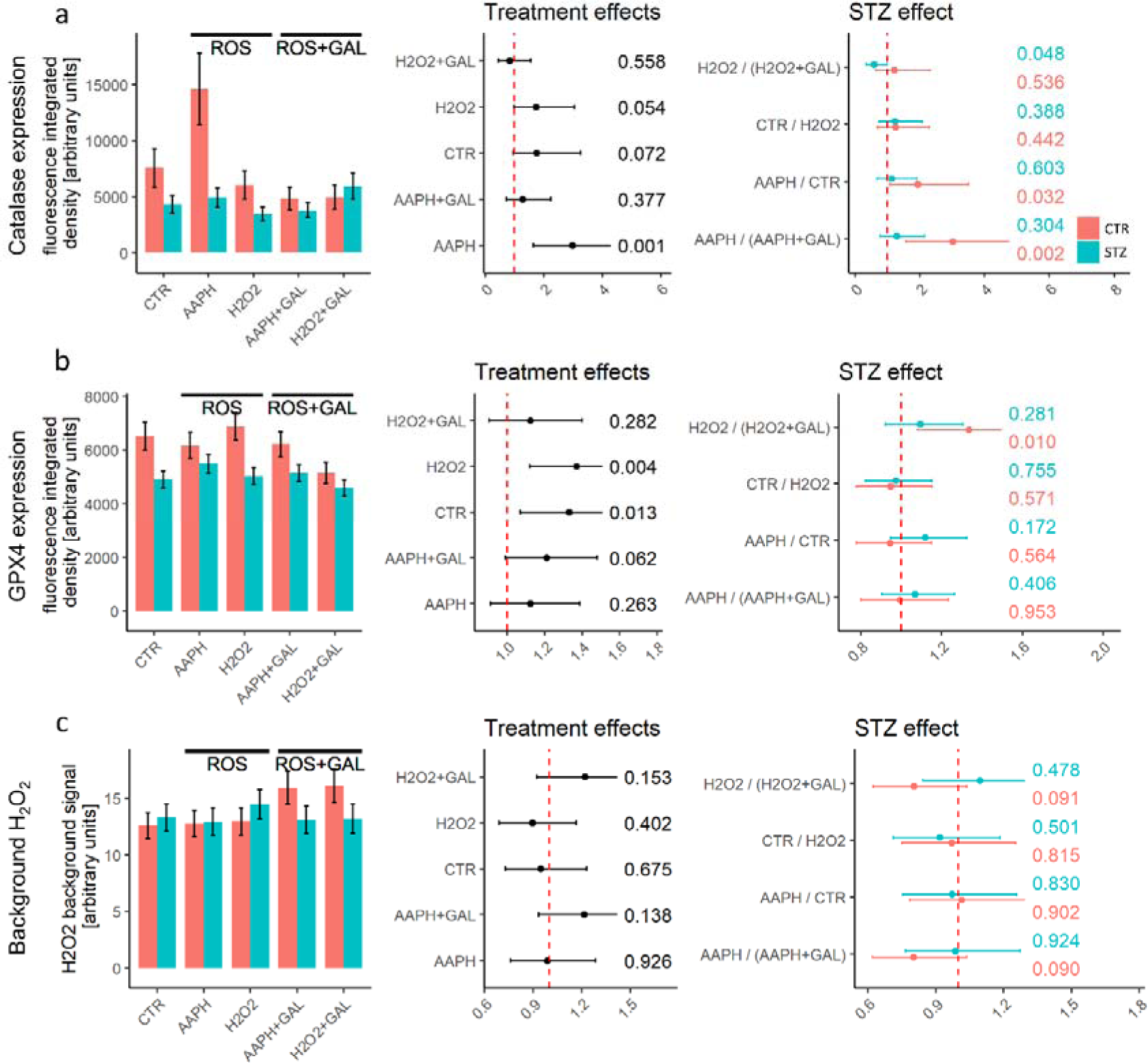
Expression of catalase (CAT) and glutathione peroxidase (GPX4) and baseline H_2_O_2_. CAT signal group estimates (left) and effect sizes (right)(a). GPX4 signal group estimates (left) and effect sizes (right)(b). Background H_2_O_2_ signal group estimates (left) and effect sizes (right)(c). The error bars in the bar graphs represent standard errors, while for effect size estimates, 95% confidence intervals are provided. P-values are reported alongside the model-derived estimates. ROS – reactive oxygen species (prooxidative environment); GAL – D-galactose.

Subsequently, we investigated the H_2_O_2_ pathway, which plays a crucial role in redox signaling. In the CTR, the presence of AAPH led to a significant induction of CAT expression, while the co-incubation with D-galactose prevented this response. In the STZ tissue, CAT expression remained relatively stable across all conditions, except when H_2_O_2_ and D-galactose were present. In this case, D-galactose prevented the H_2_O_2_-induced suppression and stimulated the expression of CAT (Fig 2a). The expression of GPX4 was consistently decreased in the STZ group across all conditions. In the control group, the presence of D-galactose resulted in reduced GPX4 expression when H_2_O_2_ was present. However, in the STZ tissue, the effects of D-galactose on GPX4 expression were absent (Fig 2B). The background H_2_O_2_ signal in the STZ group remained unchanged in all conditions, while in the controls, it increased with the presence of D-galactose (Fig 2C).

CAT activity showed no significant changes across conditions in the STZ group, indicating the reduced responsiveness of STZ cells, whereas in the control group, CAT was activated in response to the oxidative environment, and the presence of D-galactose further stimulated CAT activity in the AAPH+GAL group (Fig 3A). Interestingly, the activation of peroxidases demonstrated a similar level of responsiveness in both the CTR and STZ tissue, with the highest activation observed in the presence of D-galactose in both groups. The only distinction between the CTR and STZ was observed when the tissue was incubated with H_2_O_2_, where the activation of the peroxidase system was exclusively observed in the STZ group (Fig 3B). The absence of CAT activation and the increased activity of peroxidases in response to H_2_O_2_ exposure point to a compensatory mechanism employed by STZ tissue to maintain redox homeostasis. This suggests that the STZ tissue relies on the activation of alternative systems for removing H_2_O_2_ to counterbalance the deficient CAT activity. The total H_2_O_2_ dissociation capacity exhibited a similar pattern to CAT activation, indicating once again the diminished responsiveness of the STZ tissue, while the greatest stimulation of H_2_O_2_ removal systems was observed in the CTR tissue exposed to AAPH and D-galactose (Fig 3C). Comparative analysis using SEMs revealed significant alterations in the H_2_O_2_ signaling system in the STZ tissue (χ^2^ p=0.0002). The SEM results indicated that the H_2_O_2_ signaling in STZ tissue relied less on CAT activation and CAT expression to regulate baseline H_2_O_2_ levels. Additionally, the covariance between the activation of the H_2_O_2_ removal system and baseline H_2_O_2_ was reversed in the STZ group, suggesting a potential decrease in the capacity of protective systems and/or an increased impact of H_2_O_2_-generating systems.

**Fig 3.**
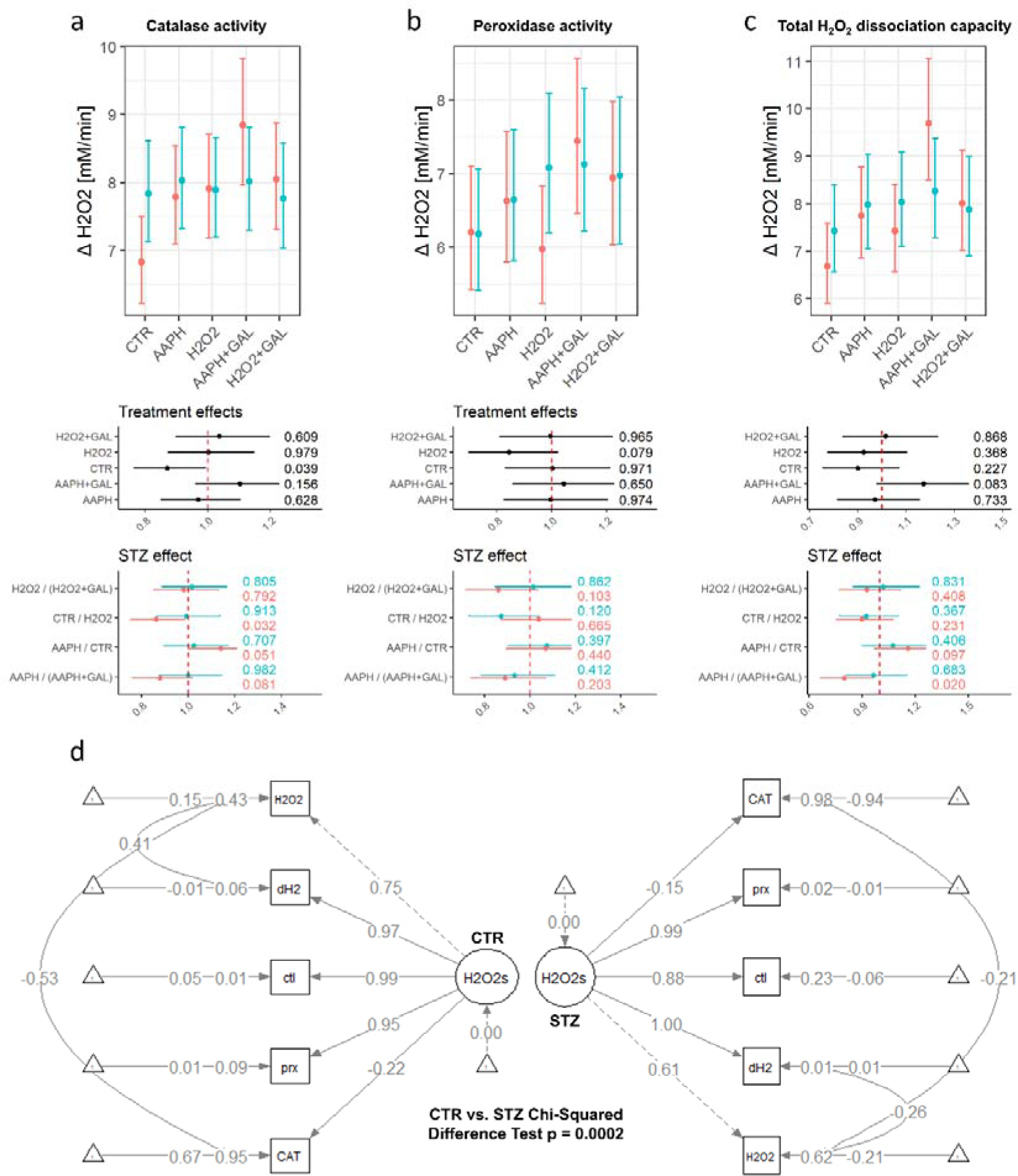
Total H_2_O_2_ dissociation capacity and the activity of catalase and peroxidases. Catalase activity group estimates (top) and effect sizes (bottom)(a). Peroxidase activity group estimates (top) and effect sizes (bottom)(b). Total H_2_O_2_ dissociation capacity group estimates (top) and effect sizes (bottom)(c). Structural equation models (SEMs) with standardized loadings for the CTR (left) and STZ (right). Manifest variables are depicted as squares, while latent variables are shown as circles. Error bars represent 95% confidence intervals are provided. P-values are reported alongside the model-derived estimates. H2O2 – background H_2_O_2_; dH2 – H_2_O_2_ dissociation capacity; ctl – catalase activity; prx – glutahione peroxidase 4 activity; CAT – catalase expression; ROS – reactive oxygen species (prooxidative environment); GAL – D-galactose.

Another important aspect of maintaining redox homeostatic control involves the regulation of O_2_^•-^ turnover and conversion into H_2_O_2_ through the SOD system. In the control group, the expression of SOD1 decreased when incubated with AAPH, whereas co-incubation with AAPH and D-galactose led to increased expression of both SOD1 and SOD2 (Fig 4A,B). The presence of H_2_O_2_ had no impact on the expression of SOD1 and SOD2, regardless of the presence of D-galactose in the control group. In contrast, the STZ samples again showed reduced responsiveness to environmental changes, as evidenced by the consistent expression levels of SOD1 and SOD2 across all conditions (Fig 4A,B).

**Fig 4.**
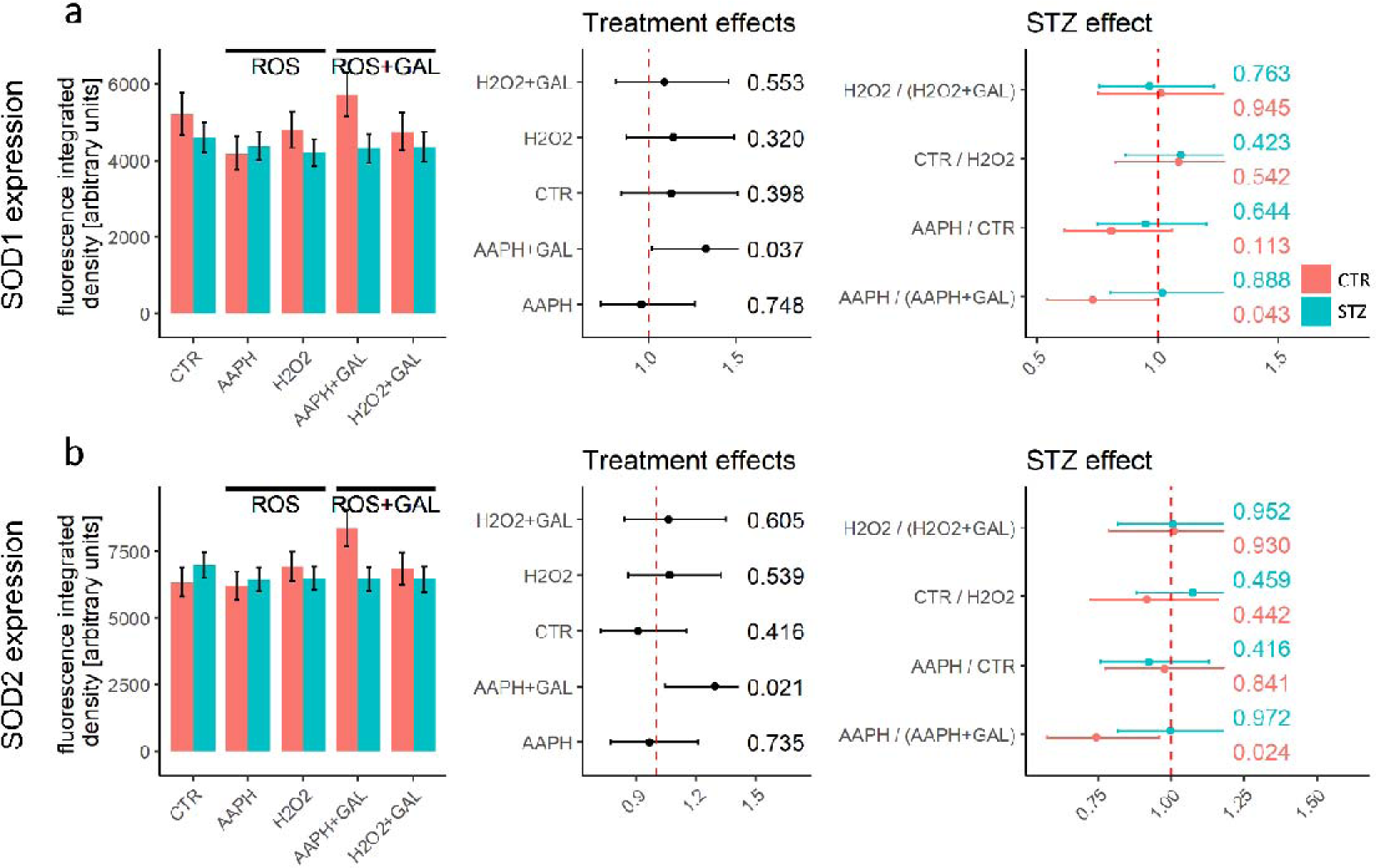
Expression of superoxide dismutase 1 (SOD1) and superoxide dismutase 2 (SOD2). SOD1 signal group estimates (left) and effect sizes (right)(a). SOD2 signal group estimates (left) and effect sizes (right)(b). The error bars in the bar graphs represent standard errors, while for effect size estimates, 95% confidence intervals are provided. P-values are reported alongside the model-derived estimates. ROS – reactive oxygen species (prooxidative environment); GAL – D-galactose.

At the baseline, SOD1 activity and total O_2_^•-^ dissociation capacity were higher in the STZ samples compared to the controls, indicating an increased demand for O_2_^•-^ removal in the former group (Fig 5A-C)). In the control group, the presence of either AAPH or H_2_O_2_ led to an increase in SOD1 activity, while SOD2 activity remained unchanged. The addition of D-galactose had no effect on SOD1 activity, SOD2 activity, or the total O_2_^•-^ dissociation capacity of the control samples. However, when D-galactose was introduced in the presence of H_2_O_2_, it reduced the activation of cytoplasmic SOD and increased the activation of mitochondrial SOD. The SEM analysis revealed a significant disturbance of the SOD system in the STZ samples (χ^2^ p=0.002). When comparing the STZ samples to the controls, it became evident that the SOD system in the former was more reliant on the activity of mitochondrial SOD, as indicated by SEM standardized estimates (Fig 5D).

**Fig 5.**
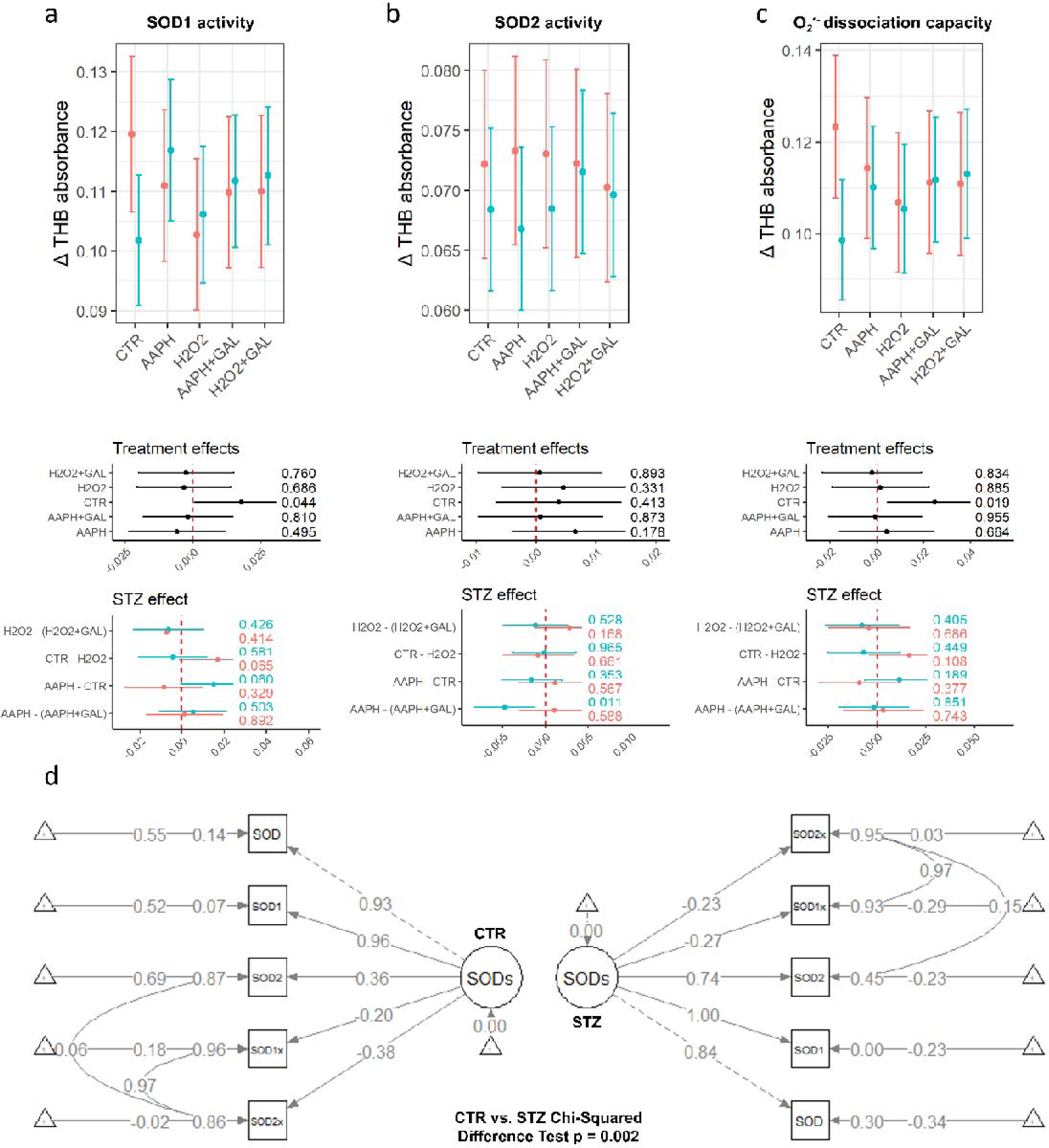
Total superoxide dissociation capacity and the activity of superoxide dismutase 1 (SOD1) and superoxide dismutase 2 (SOD2). The estimates of THB absorbance difference are inversely proportional to the removal of superoxide radicals. SOD1 activity group estimates (top) and effect sizes (bottom)(a). SOD2 activity group estimates (top) and effect sizes (bottom)(b). Total superoxide dissociation capacity group estimates (top) and effect sizes (bottom)(c). Structural equation models (SEMs) with standardized loadings for the CTR (left) and STZ (right). Manifest variables are depicted as squares, while latent variables are shown as circles. Error bars represent 95% confidence intervals are provided. P-values are reported alongside the model-derived estimates. SOD – total superoxide dissociation capacity; SOD1 – SOD1 activity; SOD2 – SOD2 activity; SOD1x – expression of SOD1; SOD2x – expression of SOD2; ROS – reactive oxygen species (prooxidative environment); GAL – D-galactose.

At the baseline, the STZ samples showed a reduction in the expression of nNOS, with a level of change similar to what could be achieved in the control samples when exposed to a prooxidative environment (Fig 6A). In the control group, the expression of nNOS decreased when the samples were incubated with AAPH or H_2_O_2_, and the addition of D-galactose had no impact on this expression. However, in the STZ samples, the expression of nNOS remained comparable across all conditions (Fig 6A). Surprisingly, despite the differences in nNOS expression, there were only minimal changes in NO levels across the various conditions in both groups (Fig 6B). Finally, lipid peroxidation levels were comparable accross groups and conditions reflecting modest changes in total antioxidant capacity biomarkers (Fig 1D,E).

**Fig 6.**
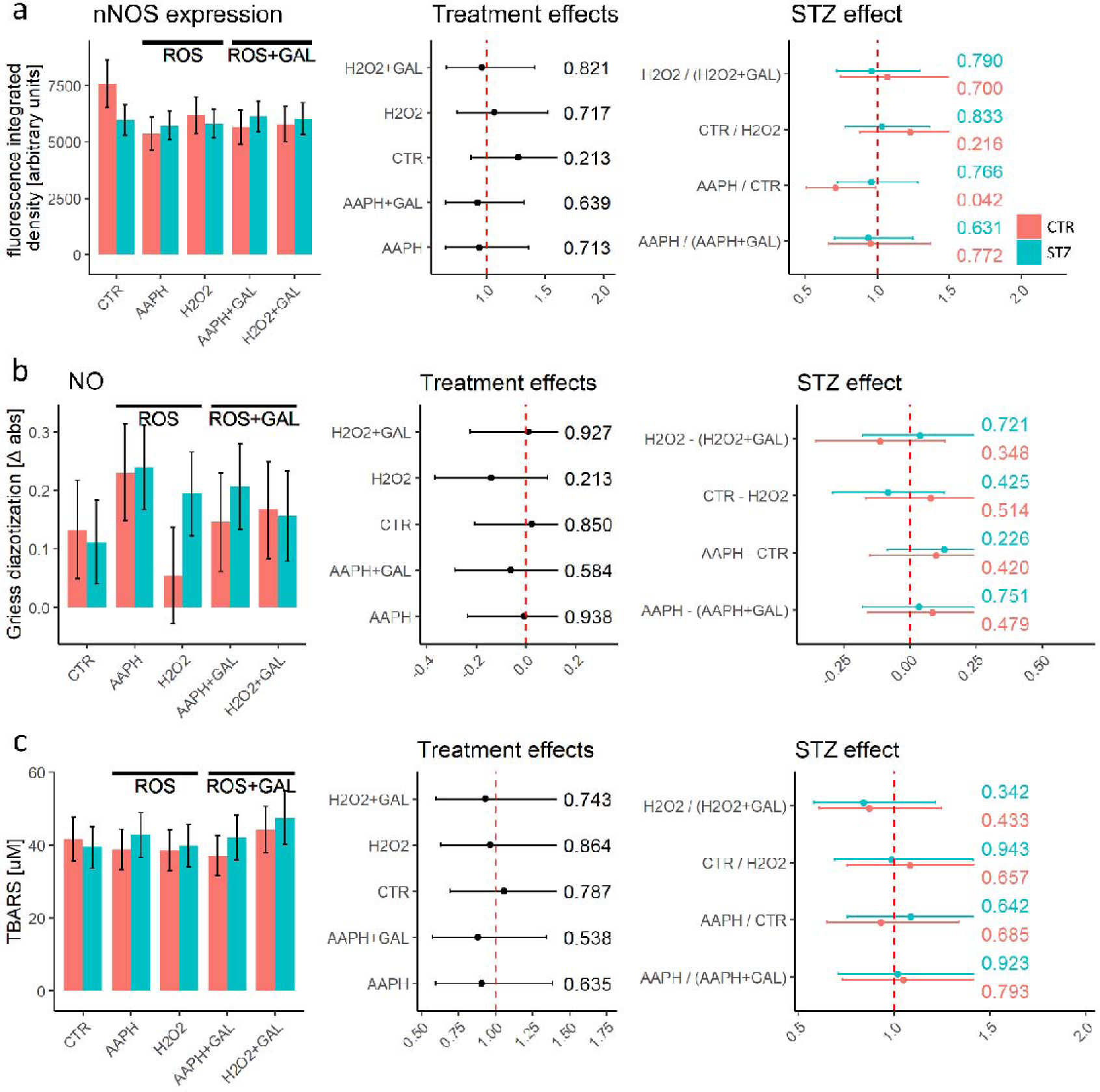
Expression of neuronal nitric oxide synthase (nNOS), quantification of nitrites and thiobarbituric acid reactive substances (TBARS). nNOS expression group estimates (left) and effect sizes (right)(a). Nitrite concentration group estimates (left) and effect sizes (right)(b). TBARS concentration group estimates (left) and effect sizes (right)(c). The error bars in the bar graphs represent standard errors, while for effect size estimates, 95% confidence intervals are provided. P-values are reported alongside the model-derived estimates. ROS – reactive oxygen species (prooxidative environment); GAL – D-galactose.

Using nonlinear dimensionality reduction through UMAP, we observed that CTR and STZ samples could be differentiated based on redox biomarkers measured in this study under normal control conditions and when exposed to prooxidative conditions, especially with AAPH (Fig 7A). However, the distinctions between the two groups became less apparent in the presence of D-galactose. This attenuation of sample dissimilarity could be attributed to the interaction between the effects of STZ and D-galactose reflected in normalization of the activity of enzymes responsible for O_2_^•-^ and H_2_O_2_ homeostasis. Another possibility is that the decrease in dissimilarity was mainly influenced by the effects of D-galactose in the CTR and its absence in the STZ group. This is indicated by the lack of distinction between the samples treated with galactose and those untreated in the STZ group (Fig 7B).

**Fig 7.**
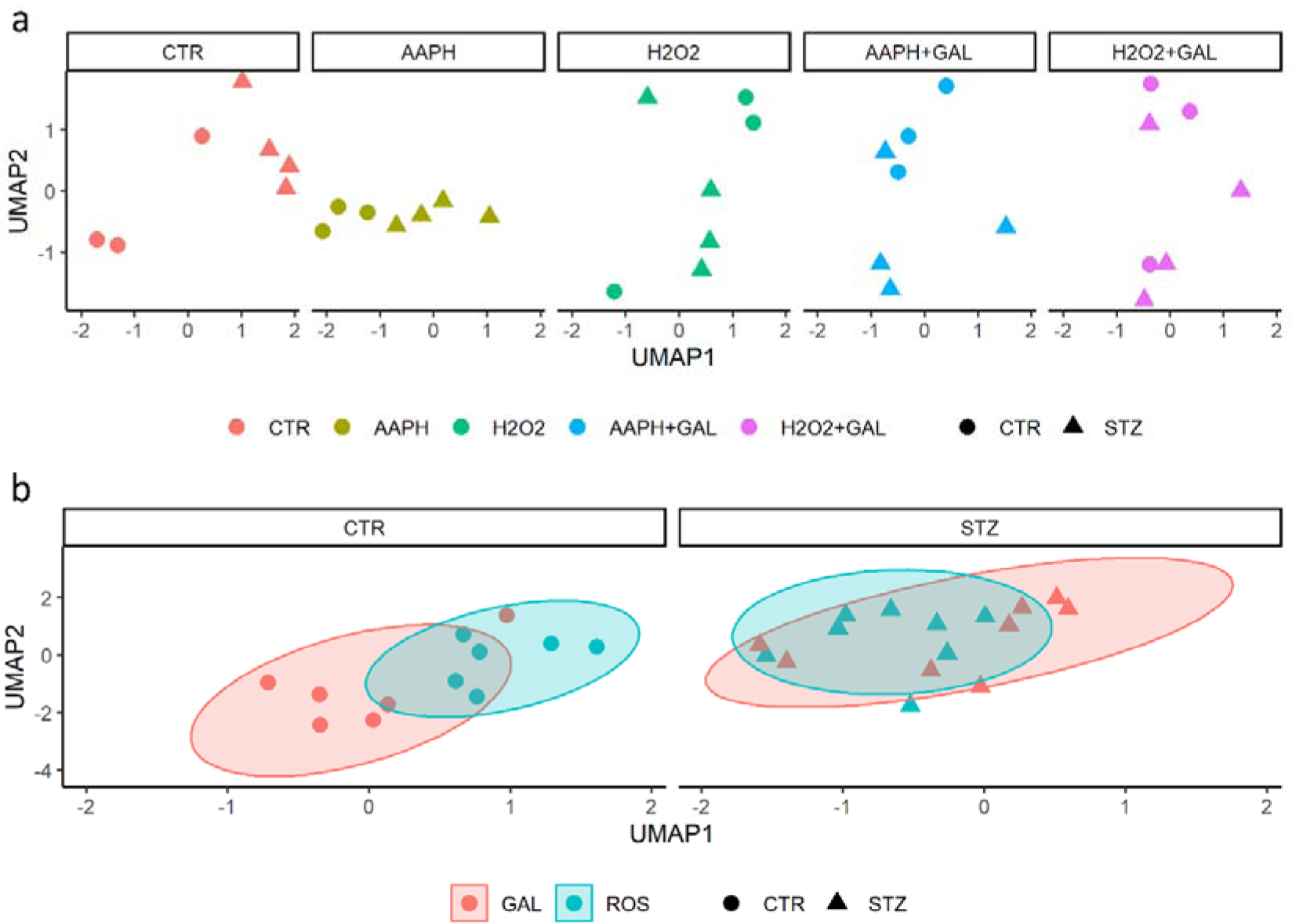
Nonlinear dimensionality reduction based on uniform manifold approximation and projection (UMAP). Positions in the UMAP biplot based on treatment conditions (a). Representation of UMAP dissimilarity of CTR and STZ samples with or without D-galactose treatament. ROS – reactive oxygen species (prooxidative environment); GAL – D-galactose.

## Discussion

The presented results offer invaluable insights into the potential mechanisms underlying the disturbance of redox balance in the gastrointestinal system of the STZ-icv rat model of AD[18]. Collectively, these findings suggest that the STZ-induced gut dysfunction is characterized by a reduced ability of the redox regulatory system to maintain long-term protection through the transcription of genes involved in antioxidant response, as well as a compromised activation of enzymes responsible for immediate antioxidant defense. Baseline expression of CAT and GPX4 showed decreased levels in the STZ samples. The tested conditions (oxidative environment; D-galactose) had no significant effect on the expression of the four redox-related proteins (CAT, SOD1, SOD2, GPX4) in STZ tissue, with one exception. Notably, CAT expression showed an increase when STZ rings were exposed to H_2_O_2_ in the presence of D-galactose. In contrast, the presence of oxidants and/or D-galactose did affect the expression of CAT, GPX4, and SODs in the control group. Due to the relatively short incubation times (30 minutes), definitive conclusions regarding protein expression cannot be drawn, and it is possible that the expression of the mentioned proteins in the STZ tissue was not entirely absent but rather delayed. Nevertheless, the data strongly suggests a lack of transcriptional/translational reactivity to oxidative stimuli and/or D-galactose in the STZ samples. The lack of redox reactivity in STZ duodenal rings was also evident in the impaired activation of enzymes crucial for immediate antioxidant defense, specifically CAT (and total H_2_O_2_ dissociation capacity), as well as SOD1 (and total O_2_^•-^ conversion capacity).

At baseline, the activity of both CAT and SOD1 was elevated in STZ tissue, suggesting heightened stress on the redox system responsible for removing O_2_^•-^ and H_2_O_2_. When exposed to oxidants, control tissue samples demonstrated a compensatory activation of CAT and SOD1, whereas duodenal rings from STZ-icv animals exhibited either absent (CAT) or reversed (SOD1) responses. The mechanisms behind the observed effects have yet to be fully understood. One possible explanation is that the O_2_^•-^ and H_2_O_2_ removal systems in the STZ gut are already functioning at their maximum activation due to a constitutively increased electrophilic tone[18]. Alternatively, the absence of response in the STZ tissue samples could be attributed to altered cellular pathways responsible for modulating the activation of antioxidant enzymes. For instance, previous research has indicated that protein kinases can swiftly regulate CAT activity[54], and the mammalian target of rapamycin (mTOR) directly phosphorylates and activates SOD1[55]. Further investigation is needed to clarify the exact mechanisms underlying these findings. In both groups, the reactivity of SOD2 activity was minimal, suggesting that its modulation was not a significant factor in the response to co-incubation with oxidants and D-galactose. However, the peroxidase system showed involvement in the response, and intriguingly, its reactivity and responsiveness were preserved in the STZ group. In fact, the total peroxidase activity showed a difference only when the samples were incubated with H_2_O_2_, suggesting that its overactivation might have been a compensatory response to the insufficient reactivity of catalase in this context. It is crucial to emphasize that the only condition in which a significant difference in the survival of mucosal cells was noticed is during incubation with H_2_O_2_ without D-galactose. The results indicate that the mucosa in the STZ group is more susceptible to H_2_O_2_-induced damage, possibly due to the reduced reactivity of the redox response. However, it is essential to consider the potential biases introduced by the decreased survival of STZ cells. First, the activity of redox-related enzymes might have been influenced by molecular signaling triggered by dying cells. For instance, in a study involving dextran sulfate sodium-induced death of colonocytes, the loss of CAT and SOD1 activity was observed[56]. A second concern is the potential introduction of attrition bias due to the loss of vulnerable cells, leaving only the least susceptible cells available for analysis. As a result, caution should be exercised when interpreting the response to H_2_O_2_ based on the reported results.

Co-incubation with D-galactose showed positive effects on redox homeostasis when cells were exposed to a prooxidative environment. However, this response was observed only in the control samples. In the control samples exposed to a prooxidative environment caused by AAPH, D-galactose enhanced the antioxidant capacity (NRP) and prevented the upregulation of CAT expression induced by AAPH. This effect may be attributed to an increase in the activity of both CAT and peroxidases, leading to approximately 45% greater total H_2_O_2_ dissociation capacity. Interestingly, while D-galactose did not impact the activity of SODs, it upregulated the expression of both SOD1 and SOD2, possibly providing long-term protection against AAPH-induced damage. When control samples were exposed to a prooxidative environment generated by H_2_O_2_, the effects on CAT activity and expression were not observed. Instead, D-galactose activated peroxidases and reduced the expression of GPX4, suggesting activation of an alternative adaptive response. Together, the results indicate that D-galactose has the capacity to positively impact gut redox homeostasis. These findings are consistent with an *in vivo* study conducted on rats, where the acute oral administration of D-galactose showed beneficial effects on the redox homeostasis of the small intestine[40]. Furthermore, these results provide additional evidence supporting the notion that D-galactose’s protective effects on the gut, such as protection against ionizing radiation[57], may be attributed to its ability to modulate redox homeostasis[58].

The STZ samples showed attenuated response to D-galactose treatment. The only noticeable effect of D-galactose in the STZ duodenal rings was an increase in mucosal cell survival, coupled with an upregulation of CAT expression when the samples were exposed to H_2_O_2_. The lack of response of STZ samples to D-galactose in the prooxidative environment is evident from the absence of spatial distinction in UMAP. The mechanistic explanation for the lack of response of STZ samples to D-galactose treatment remains elusive. However, previous experiments have indicated that the beneficial effects of D-galactose on redox homeostasis rely on the utilization of reductive equivalents, such as low-molecular-weight thiols (LMWT) and reduced nicotinamide adenine dinucleotide phosphate (NADPH)[40], possibly due to its hormetic effects[41]. This suggests that the reduced concentration of LMWT in the duodenum, as observed in the STZ-icv model of AD[18], could potentially contribute to the absence of a positive response to D-galactose treatment observed in this study.

Previous experiments have revealed that duodenal goblet cells in the STZ-icv rat model of AD do not respond to cholinergic stimulation for mucus secretion [20]. This study builds upon those findings by demonstrating that the GI tract in STZ-icv rats also exhibits reduced responsiveness to an oxidative environment and exposure to D-galactose. Consequently, this suggests that the GI tract of STZ-icv rats may generally be considered unresponsive to both internal and external stimuli.

To gain a deeper understanding of the underlying pathophysiological processes in the gut resulting from central neurodegeneration, future experiments should focus on investigating how changes in the CNS contribute to the observed unresponsive behavior in GI cells. This aspect is particularly intriguing considering that redox dyshomeostasis and the “unresponsive” gut phenotype have not been observed in other brain-first toxin-induced models of neurodegenerative diseases. This indicates that the damage induced by STZ-icv likely does not solely affect the GI tract by damaging the efferent control of gut function, such as via the efferent arm of the vagus nerve.

For instance, in the context of mimicking Parkinson’s disease through intrastriatal administration of 6-hydroxydopamine, central neurodegeneration does not seem to lead to pronounced redox dyshomeostasis in the gut [21]. Although GI changes have been observed in this model [59–61], they cannot be solely attributed to the absence of responsiveness to stimuli as a common factor.

## Conclusion

Together, the presented findings offer crucial insights into the mechanisms of redox dyshomeostasis in the gastrointestinal (GI) tract of the STZ-icv rat model of Alzheimer’s disease (AD), as well as important information regarding the protective effects of D-galactose in the gut. In the STZ-induced gut dysfunction, there is a noticeable reduction in the ability of the redox regulatory system to maintain protection against free radicals, evidenced by the decreased activation of enzymes responsible for immediate antioxidant defense and the upregulation of genes involved in the antioxidant response. The results obtained from control samples exposed to a prooxidative environment in the presence of D-galactose indicate that D-galactose can have beneficial effects on redox homeostasis. Specifically, it enhances antioxidant capacity, increases the activity of CAT and peroxidases, and stimulates the expression of SODs. However, these advantageous effects of D-galactose on redox homeostasis are less pronounced in the STZ-icv model of AD.

### Limitations

The presented study has several important limitations that warrant consideration. Firstly, although *ex vivo* experiments provide valuable insights into mechanisms by enabling precise control over experimental conditions, they may not entirely replicate the complex molecular events occurring in the actual *in vivo* environment. It’s crucial to acknowledge this potential discrepancy when interpreting the results. Secondly, in this study, prooxidant conditions were simulated by subjecting duodenal rings to a single concentration of AAPH and H_2_O_2_, which aligns with established standards in the literature [62,63]. Moreover, to minimize bias introduced by decreased cell survival, the reaction was halted at a single time-point (after 30 minutes of incubation). However, it’s important to recognize that varying exposure times and concentrations of AAPH and H_2_O_2_ could lead to different outcomes and conclusions, and these alternative conditions should be considered in future investigations. Additionally, although we assessed cell survival in duodenal sections, we were unable to determine the spatial distribution of the expression of redox-related enzymes due to limited availability of tissue samples. Consequently, we cannot pinpoint the specific cells driving the observed changes at the level of whole tissue homogenate. While post-treatment microdissection was contemplated, the decreased survival of cells during the dissection process was associated with experimental error, leading us to dismiss this approach to avoid introducing bias. Therefore, it is essential to take these limitations into account when interpreting the study results and consider future research efforts to address these aspects comprehensively.

## Author contributions

JH – study design; ABP, AK, DV, JOB, JH – model induction, behavioral assessment; JH – ex vivo experiment; KV, VS, JH – biochemical analyses; KV, VS – tissue processing, image acquisition; JH, KV – data curation; JH – data analysis, writing the first draft of the manuscript. KV, VS, ABP, AK, DV, JOB, MSP – critical revision of the manuscript. MSP – funding, supervision.

## Data Availability

Raw data for the study can be obtained from the corresponding author upon request. Preprinted on bioRxiv.

## Ethics approval

The animal procedures were conducted following the guidelines of the University of Zagreb School of Medicine, as well as the national regulations outlined in The Animal Protection Act (NN135/2006; NN 47/2011) and international guidelines from Directive 2010/63/EU on the use of experimental animals. Approval for the experiments was obtained from both the Croatian Ministry of Agriculture (EP 186 /2018; 525-10/0255-15-5) and the Ethical Committee of the University of Zagreb School of Medicine (380-59-10106-18-111/173).

## Consent to participate

Not applicable.

## Consent for publication

Not applicable.

## Acknowledgments

None.

## References

[1] A. Singh, T.M. Dawson, S. Kulkarni, Neurodegenerative disorders and gut-brain interactions, J Clin Invest. 131 (2021) e143775. https://doi.org/10.1172/JCI143775.

[2] A. Bhattacharyya, R. Chattopadhyay, S. Mitra, S.E. Crowe, Oxidative stress: an essential factor in the pathogenesis of gastrointestinal mucosal diseases, Physiol Rev. 94 (2014) 329–354. https://doi.org/10.1152/physrev.00040.2012.

[3] S.C. Bischoff, G. Barbara, W. Buurman, T. Ockhuizen, J.-D. Schulzke, M. Serino, H. Tilg, A. Watson, J.M. Wells, Intestinal permeability--a new target for disease prevention and therapy, BMC Gastroenterol. 14 (2014) 189. https://doi.org/10.1186/s12876-014-0189-7.

[4] J. Homolak, Gastrointestinal redox homeostasis in ageing, Biogerontology. (2023). https://doi.org/10.1007/s10522-023-10049-8.

[5] J. Homolak, Targeting the microbiota-mitochondria crosstalk in neurodegeneration with senotherapeutics, Adv Protein Chem Struct Biol. 136 (2023) 339–383. https://doi.org/10.1016/bs.apcsb.2023.02.018.

[6] B. Chassaing, M. Kumar, M.T. Baker, V. Singh, M. Vijay-Kumar, Mammalian gut immunity, Biomed J. 37 (2014) 246–258. https://doi.org/10.4103/2319-4170.130922.

[7] S.S. Alves, R.M.P. da Silva-Junior, G. Servilha-Menezes, J. Homolak, M. Šalković-Petrišić, N. Garcia-Cairasco, Insulin Resistance as a Common Link Between Current Alzheimer’s Disease Hypotheses, J Alzheimers Dis. 82 (2021) 71–105. https://doi.org/10.3233/JAD-210234.

[8] J. Homolak, Redox Homeostasis in Alzheimer’s Disease, in: U. Çakatay (Ed.), Redox Signaling and Biomarkers in Ageing, Springer International Publishing, Cham, 2022: pp. 323–348. https://doi.org/10.1007/978-3-030-84965-8_15.

[9] K.J. Barnham, C.L. Masters, A.I. Bush, Neurodegenerative diseases and oxidative stress, Nat Rev Drug Discov. 3 (2004) 205–214. https://doi.org/10.1038/nrd1330.

[10] V.H. Perry, C. Cunningham, C. Holmes, Systemic infections and inflammation affect chronic neurodegeneration, Nat Rev Immunol. 7 (2007) 161–167. https://doi.org/10.1038/nri2015.

[11] G.C. Brown, The endotoxin hypothesis of neurodegeneration, J Neuroinflammation. 16 (2019) 180. https://doi.org/10.1186/s12974-019-1564-7.

[12] X.-H. Qian, X.-X. Song, X.-L. Liu, S. Chen, H.-D. Tang, Inflammatory pathways in Alzheimer’s disease mediated by gut microbiota, Ageing Res Rev. 68 (2021) 101317. https://doi.org/10.1016/j.arr.2021.101317.

[13] M. Carabotti, A. Scirocco, M.A. Maselli, C. Severi, The gut-brain axis: interactions between enteric microbiota, central and enteric nervous systems, Ann Gastroenterol. 28 (2015) 203– 209. https://www.ncbi.nlm.nih.gov/pmc/articles/PMC4367209/ (accessed April 1, 2023).

[14] M. Herath, S. Hosie, J.C. Bornstein, A.E. Franks, E.L. Hill-Yardin, The Role of the Gastrointestinal Mucus System in Intestinal Homeostasis: Implications for Neurological Disorders, Front Cell Infect Microbiol. 10 (2020) 248. https://doi.org/10.3389/fcimb.2020.00248.

[15] P. Honarpisheh, C.R. Reynolds, M.P. Blasco Conesa, J.F. Moruno Manchon, N. Putluri, M.B. Bhattacharjee, A. Urayama, L.D. McCullough, B.P. Ganesh, Dysregulated Gut Homeostasis Observed Prior to the Accumulation of the Brain Amyloid-β in Tg2576 Mice, Int J Mol Sci. 21 (2020) 1711. https://doi.org/10.3390/ijms21051711.

[16] S. Semar, M. Klotz, M. Letiembre, C. Van Ginneken, A. Braun, V. Jost, M. Bischof, W.J. Lammers, Y. Liu, K. Fassbender, T. Wyss-Coray, F. Kirchhoff, K.-H. Schäfer, Changes of the enteric nervous system in amyloid-β protein precursor transgenic mice correlate with disease progression, J Alzheimers Dis. 36 (2013) 7–20. https://doi.org/10.3233/JAD-120511.

[17] J. Homolak, A.B. Perhoc, A. Knezovic, J.O. Barilar, D. Virag, M. Salkovic-Petrisic, An exploratory study of gastrointestinal redox biomarkers in the presymptomatic and symptomatic Tg2576 mouse model of familial Alzheimer’s disease – phenotypic correlates and the effects of chronic oral D-galactose, bioRxiv. (2023) 2023.06.03.542513. https://doi.org/10.1101/2023.06.03.542513.

[18] J. Homolak, A. Babic Perhoc, A. Knezovic, J. Osmanovic Barilar, M. Salkovic-Petrisic, Failure of the Brain Glucagon-Like Peptide-1-Mediated Control of Intestinal Redox Homeostasis in a Rat Model of Sporadic Alzheimer’s Disease, Antioxidants (Basel). 10 (2021) 1118. https://doi.org/10.3390/antiox10071118.

[19] J. Homolak, Pathophysiological alterations of gastrointestinal system in animal models of Alzheimer’s and Parkinson’s disease, info:eu-repo/semantics/doctoralThesis, University of Zagreb. School of Medicine, 2023. https://urn.nsk.hr/urn:nbn:hr:105:754088 (accessed July 1, 2023).

[20] J. Homolak, J. De Busscher, M. Zambrano-Lucio, M. Joja, D. Virag, A. Babic Perhoc, A. Knezovic, J. Osmanovic Barilar, M. Salkovic-Petrisic, Altered Secretion, Constitution, and Functional Properties of the Gastrointestinal Mucus in a Rat Model of Sporadic Alzheimer’s Disease, ACS Chem Neurosci. (2023). https://doi.org/10.1021/acschemneuro.3c00223.

[21] J. Homolak, M. Joja, G. Grabaric, E. Schiatti, D. Virag, A.B. Perhoc, A. Knezovic, J.O. Barilar, M. Salkovic-Petrisic, The absence of gastrointestinal redox dyshomeostasis in the brain-first rat model of Parkinson’s disease induced by bilateral intrastriatal 6-hydroxydopamine, BioRxiv. (2022) 2022.08.22.504759. https://doi.org/10.1101/2022.08.22.504759.

[22] M. Salkovic-Petrisic, A.B. Perhoc, J. Homolak, A. Knezovic, J. Osmanovic Barilar, P. Riederer, Experimental Approach to Alzheimer’s Disease with Emphasis on Insulin Resistance in the Brain, in: R.M. Kostrzewa (Ed.), Handbook of Neurotoxicity, Springer International Publishing, Cham, 2021: pp. 1–52. https://doi.org/10.1007/978-3-030-71519-9_98-1.

[23] J. Homolak, A.B. Perhoc, A. Knezovic, J. Osmanovic Barilar, M. Salkovic-Petrisic, Additional methodological considerations regarding optimization of the dose of intracerebroventricular streptozotocin A response to: “Optimization of intracerebroventricular streptozotocin dose for the induction of neuroinflammation and memory impairments in rats” by Ghosh et al., Metab Brain Dis 2020 July 21, Metab Brain Dis. 36 (2021) 97–102. https://doi.org/10.1007/s11011-020-00637-9.

[24] S.C. Correia, R.X. Santos, G. Perry, X. Zhu, P.I. Moreira, M.A. Smith, Insulin-Resistant Brain State: the culprit in sporadic Alzheimer’s Disease?, Ageing Res Rev. 10 (2011) 264–273. https://doi.org/10.1016/j.arr.2011.01.001.

[25] A. Knezovic, J. Osmanovic Barilar, A. Babic, R. Bagaric, V. Farkas, P. Riederer, M. Salkovic-Petrisic, Glucagon-like peptide-1 mediates effects of oral galactose in streptozotocin-induced rat model of sporadic Alzheimer’s disease, Neuropharmacology. 135 (2018) 48–62. https://doi.org/10.1016/j.neuropharm.2018.02.027.

[26] M. Sharma, Y.K. Gupta, Intracerebroventricular injection of streptozotocin in rats produces both oxidative stress in the brain and cognitive impairment, Life Sci. 68 (2001) 1021–1029. https://doi.org/10.1016/s0024-3205(00)01005-5.

[27] A. Knezovic, A. Loncar, J. Homolak, U. Smailovic, J. Osmanovic Barilar, L. Ganoci, N. Bozina, P. Riederer, M. Salkovic-Petrisic, Rat brain glucose transporter-2, insulin receptor and glial expression are acute targets of intracerebroventricular streptozotocin: risk factors for sporadic Alzheimer’s disease?, J Neural Transm (Vienna). 124 (2017) 695–708. https://doi.org/10.1007/s00702-017-1727-6.

[28] R. Ghosh, S. Sil, P. Gupta, T. Ghosh, Optimization of intracerebroventricular streptozotocin dose for the induction of neuroinflammation and memory impairments in rats, Metab Brain Dis. 35 (2020) 1279–1286. https://doi.org/10.1007/s11011-020-00588-1.

[29] Y. Li, P. Xu, J. Shan, W. Sun, X. Ji, T. Chi, P. Liu, L. Zou, Interaction between hyperphosphorylated tau and pyroptosis in forskolin and streptozotocin induced AD models, Biomed Pharmacother. 121 (2020) 109618. https://doi.org/10.1016/j.biopha.2019.109618.

[30] M. Salkovic-Petrisic, J. Osmanovic-Barilar, M.K. Brückner, S. Hoyer, T. Arendt, P. Riederer, Cerebral amyloid angiopathy in streptozotocin rat model of sporadic Alzheimer’s disease: a long-term follow up study, J Neural Transm (Vienna). 118 (2011) 765–772. https://doi.org/10.1007/s00702-011-0651-4.

[31] E. Grünblatt, J. Homolak, A. Babic Perhoc, V. Davor, A. Knezovic, J. Osmanovic Barilar, P. Riederer, S. Walitza, C. Tackenberg, M. Salkovic-Petrisic, From attention-deficit hyperactivity disorder to sporadic Alzheimer’s disease-Wnt/mTOR pathways hypothesis, Front Neurosci. 17 (2023) 1104985. https://doi.org/10.3389/fnins.2023.1104985.

[32] A. Knezovic, J. Osmanovic-Barilar, M. Curlin, P.R. Hof, G. Simic, P. Riederer, M. Salkovic-Petrisic, Staging of cognitive deficits and neuropathological and ultrastructural changes in streptozotocin-induced rat model of Alzheimer’s disease, J Neural Transm (Vienna). 122 (2015) 577–592. https://doi.org/10.1007/s00702-015-1394-4.

[33] J. Homolak, A. Babic Perhoc, A. Knezovic, J. Osmanovic Barilar, F. Koc, C. Stanton, R.P. Ross, M. Salkovic-Petrisic, Disbalance of the Duodenal Epithelial Cell Turnover and Apoptosis Accompanies Insensitivity of Intestinal Redox Homeostasis to Inhibition of the Brain Glucose-Dependent Insulinotropic Polypeptide Receptors in a Rat Model of Sporadic Alzheimer’s Disease, Neuroendocrinology. 112 (2022) 744–762. https://doi.org/10.1159/000519988.

[34] J. Osmanović Barilar, A. Babić Perhoč, A. Knezović, J. Homolak, D. Virag, M. Šalković-Petrišić, The Effect of the Sodium-Glucose Cotransporter Inhibitor on Cognition and Metabolic Parameters in a Rat Model of Sporadic Alzheimer’s Disease, Biomedicines. 11 (2023) 1025. https://doi.org/10.3390/biomedicines11041025.

[35] R.C.M. de Moraes, G.C.A. Lima, C.A.E.F. Cardinali, A.C. Gonçalves, G.V. Portari, E.M. Guerra-Shinohara, A. Leboucher, J. Donato, A. Kleinridders, A. da S. Torrão, Benfotiamine protects against hypothalamic dysfunction in a STZ-induced model of neurodegeneration in rats, Life Sci. 306 (2022) 120841. https://doi.org/10.1016/j.lfs.2022.120841.

[36] S. Sadigh-Eteghad, A. Majdi, S.K. McCann, J. Mahmoudi, M.S. Vafaee, M.R. Macleod, D-galactose-induced brain ageing model: A systematic review and meta-analysis on cognitive outcomes and oxidative stress indices, PLoS One. 12 (2017) e0184122. https://doi.org/10.1371/journal.pone.0184122.

[37] A. Babic Perhoc, J. Osmanovic Barilar, A. Knezovic, V. Farkas, R. Bagaric, A. Svarc, E. Grünblatt, P. Riederer, M. Salkovic-Petrisic, Cognitive, behavioral and metabolic effects of oral galactose treatment in the transgenic Tg2576 mice, Neuropharmacology. 148 (2019) 50–67. https://doi.org/10.1016/j.neuropharm.2018.12.018.

[38] M. Salkovic-Petrisic, Oral Galactose Provides a Different Approach to Incretin-Based Therapy of Alzheimer’s Disease, Journal of Neurology & Neuromedicine. 3 (2018). https://www.jneurology.com/articles/oral-galactose-provides-a-different-approach-to-incretinbased-therapy-of-alzheimers-disease.html (accessed March 25, 2023).

[39] M. Salkovic-Petrisic, J. Osmanovic-Barilar, A. Knezovic, S. Hoyer, K. Mosetter, W. Reutter, Long-term oral galactose treatment prevents cognitive deficits in male Wistar rats treated intracerebroventricularly with streptozotocin, Neuropharmacology. 77 (2014) 68–80. https://doi.org/10.1016/j.neuropharm.2013.09.002.

[40] J. Homolak, A. Babic Perhoc, A. Knezovic, J. Osmanovic Barilar, D. Virag, M. Joja, M. Salkovic-Petrisic, The Effect of Acute Oral Galactose Administration on the Redox System of the Rat Small Intestine, Antioxidants (Basel). 11 (2021) 37. https://doi.org/10.3390/antiox11010037.

[41] J. Homolak, A. Babic Perhoc, A. Knezovic, I. Kodvanj, D. Virag, J. Osmanovic Barilar, P. Riederer, M. Salkovic-Petrisic, Is Galactose a Hormetic Sugar? An Exploratory Study of the Rat Hippocampal Redox Regulatory Network, Mol Nutr Food Res. 65 (2021) e2100400. https://doi.org/10.1002/mnfr.202100400.

[42] E.P. Noble, R.J. Wurtman, J. Axelrod, A simple and rapid method for injecting H3-norepinephrine into the lateral ventricle of the rat brain, Life Sci. 6 (1967) 281–291. https://doi.org/10.1016/0024-3205(67)90157-9.

[43] I.R. Ilyasov, V.L. Beloborodov, I.A. Selivanova, R.P. Terekhov, ABTS/PP Decolorization Assay of Antioxidant Capacity Reaction Pathways, Int J Mol Sci. 21 (2020) 1131. https://doi.org/10.3390/ijms21031131.

[44] J. Homolak, I. Kodvanj, A. Babic Perhoc, D. Virag, A. Knezovic, J. Osmanovic Barilar, P. Riederer, M. Salkovic-Petrisic, Nitrocellulose redox permanganometry: A simple method for reductive capacity assessment, MethodsX. 9 (2022) 101611. https://doi.org/10.1016/j.mex.2021.101611.

[45] E. Hrabárová, K. Valachová, P. Rapta, L. Soltés, An alternative standard for Trolox-equivalent antioxidant-capacity estimation based on thiol antioxidants. Comparative 2,2’-azinobis[3-ethylbenzothiazoline-6-sulfonic acid] decolorization and rotational viscometry study regarding hyaluronan degradation, Chem Biodivers. 7 (2010) 2191–2200. https://doi.org/10.1002/cbdv.201000019.

[46] S. Marklund, G. Marklund, Involvement of the superoxide anion radical in the autoxidation of pyrogallol and a convenient assay for superoxide dismutase, Eur J Biochem. 47 (1974) 469–474. https://doi.org/10.1111/j.1432-1033.1974.tb03714.x.

[47] X. Li, Improved pyrogallol autoxidation method: a reliable and cheap superoxide-scavenging assay suitable for all antioxidants, J Agric Food Chem. 60 (2012) 6418–6424. https://doi.org/10.1021/jf204970r.

[48] J. Homolak, In vitro analysis of catalase and superoxide dismutase mimetic properties of blue tattoo ink, Free Radic Res. 56 (2022) 343–357. https://doi.org/10.1080/10715762.2022.2102976.

[49] M.H. Hadwan, Simple spectrophotometric assay for measuring catalase activity in biological tissues, BMC Biochem. 19 (2018) 7. https://doi.org/10.1186/s12858-018-0097-5.

[50] J. Homolak, The effect of a color tattoo on the local skin redox regulatory network: an N-of-1 study, Free Radic Res. 55 (2021) 221–229. https://doi.org/10.1080/10715762.2021.1912340.

[51] X. Ma, D. Deng, W. Chen, X. Ma, D. Deng, W. Chen, Inhibitors and Activators of SOD, GSH-Px, and CAT, IntechOpen, 2017. https://doi.org/10.5772/65936.

[52] N. Percie du Sert, A. Ahluwalia, S. Alam, M.T. Avey, M. Baker, W.J. Browne, A. Clark, I.C. Cuthill, U. Dirnagl, M. Emerson, P. Garner, S.T. Holgate, D.W. Howells, V. Hurst, N.A. Karp, S.E. Lazic, K. Lidster, C.J. MacCallum, M. Macleod, E.J. Pearl, O.H. Petersen, F. Rawle, P. Reynolds, K. Rooney, E.S. Sena, S.D. Silberberg, T. Steckler, H. Würbel, Reporting animal research: Explanation and elaboration for the ARRIVE guidelines 2.0, PLoS Biol. 18 (2020) e3000411. https://doi.org/10.1371/journal.pbio.3000411.

[53] Y. Rosseel, lavaan: An R Package for Structural Equation Modeling, Journal of Statistical Software. 48 (2012) 1–36. https://doi.org/10.18637/jss.v048.i02.

[54] S. Yano, N. Yano, Regulation of catalase enzyme activity by cell signaling molecules, Mol Cell Biochem. 240 (2002) 119–130. https://doi.org/10.1023/a:1020680131754.

[55] C.K. Tsang, M. Chen, X. Cheng, Y. Qi, Y. Chen, I. Das, X. Li, B. Vallat, L.-W. Fu, C.-N. Qian, H.-Y. Wang, E. White, S.K. Burley, X.F.S. Zheng, SOD1 Phosphorylation by mTORC1 Couples Nutrient Sensing and Redox Regulation, Mol Cell. 70 (2018) 502–515.e8. https://doi.org/10.1016/j.molcel.2018.03.029.

[56] J. Hwang, J. Jin, S. Jeon, S.H. Moon, M.Y. Park, D.-Y. Yum, J.H. Kim, J.-E. Kang, M.H. Park, E.-J. Kim, J.-G. Pan, O. Kwon, G.T. Oh, SOD1 suppresses pro-inflammatory immune responses by protecting against oxidative stress in colitis, Redox Biol. 37 (2020) 101760. https://doi.org/10.1016/j.redox.2020.101760.

[57] T. Zhu, Z. Wang, J. He, X. Zhang, C. Zhu, S. Zhang, Y. Li, S. Fan, D-galactose protects the intestine from ionizing radiation-induced injury by altering the gut microbiome, J Radiat Res. 63 (2022) 805–816. https://doi.org/10.1093/jrr/rrac059.

[58] J. Homolak, A. Babic Perhoc, D. Virag, A. Knezovic, J. Osmanovic Barilar, M. Salkovic-Petrisic, D-galactose might protect against ionizing radiation by stimulating oxidative metabolism and modulating redox homeostasis, J Radiat Res. 64 (2023) 743–745. https://doi.org/10.1093/jrr/rrad046.

[59] M. Colucci, M. Cervio, M. Faniglione, S. De Angelis, M. Pajoro, G. Levandis, C. Tassorelli, F. Blandini, F. Feletti, R. De Giorgio, A. Dellabianca, S. Tonini, M. Tonini, Intestinal dysmotility and enteric neurochemical changes in a Parkinson’s disease rat model, Auton Neurosci. 169 (2012) 77–86. https://doi.org/10.1016/j.autneu.2012.04.005.

[60] H. Cui, J.D. Elford, O. Alitalo, P. Perez-Pardo, J. Tampio, K.M. Huttunen, A. Kraneveld, M.M. Forsberg, T.T. Myöhänen, A.J. Jalkanen, Nigrostriatal 6-hydroxydopamine lesions increase alpha-synuclein levels and permeability in rat colon, Neurobiol Aging. 129 (2023) 62–71. https://doi.org/10.1016/j.neurobiolaging.2023.05.007.

[61] H.C. Zhu, J. Zhao, C.Y. Luo, Q.Q. Li, Gastrointestinal dysfunction in a Parkinson’s disease rat model and the changes of dopaminergic, nitric oxidergic, and cholinergic neurotransmitters in myenteric plexus, J Mol Neurosci. 47 (2012) 15–25. https://doi.org/10.1007/s12031-011-9560-0.

[62] Y. Dong, Q. Hou, J. Lei, P.G. Wolf, H. Ayansola, B. Zhang, Quercetin Alleviates Intestinal Oxidative Damage Induced by H2O2 via Modulation of GSH: In Vitro Screening and In Vivo Evaluation in a Colitis Model of Mice, ACS Omega. 5 (2020) 8334–8346. https://doi.org/10.1021/acsomega.0c00804.

[63] J. Wu, B. Sun, X. Luo, M. Zhao, F. Zheng, J. Sun, H. Li, X. Sun, M. Huang, Cytoprotective effects of a tripeptide from Chinese Baijiu against AAPH-induced oxidative stress in HepG2 cells via Nrf2 signaling, RSC Adv. 8 (2018) 10898–10906. https://doi.org/10.1039/c8ra01162a.

